## Supplementary figures and tables for "Citalopram exhibits immune-dependent anti-tumor effects by modulating C5aR1^+^ TAMs"

### Supplementary Materials

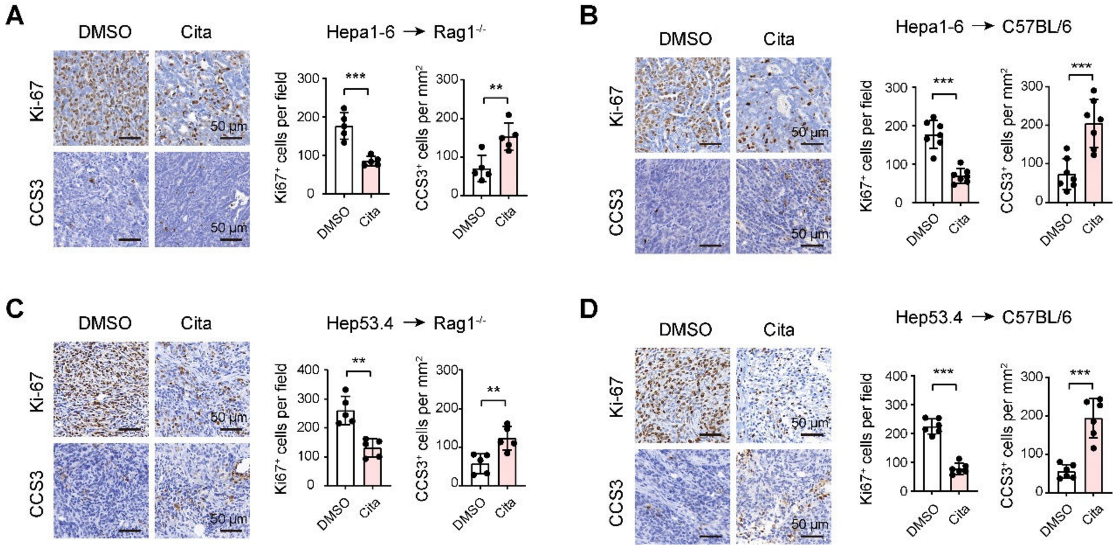


**Figure S1. Citalopram inhibits HCC cell proliferation and promotes cell apoptosis in the immune-competent and immune-deficient mouse models.**

(**A**, **B**) Immunohistochemical analysis of cleaved caspase-3 (CCS3) and Ki67 in Hepa1-6-bearing subcutaneous xenograft tumors from $Rag1^{-/-}$ or immunocompetent C57BL/6 mice, treated with DMSO or 5 mg/kg citalopram. (**C**, **D**) Immunohistochemical analysis of CCS3 and Ki67 in Hep53.4-bearing subcutaneous xenograft tumors from $Rag1^{-/-}$ or immunocompetent C57BL/6 mice, treated with DMSO or 5 mg/kg citalopram. In all panels, $^{**}p < 0.01$, $^{***}p < 0.001$. Scale bar, 50 μm. Values as mean ± SD and compared by the Student's t test.

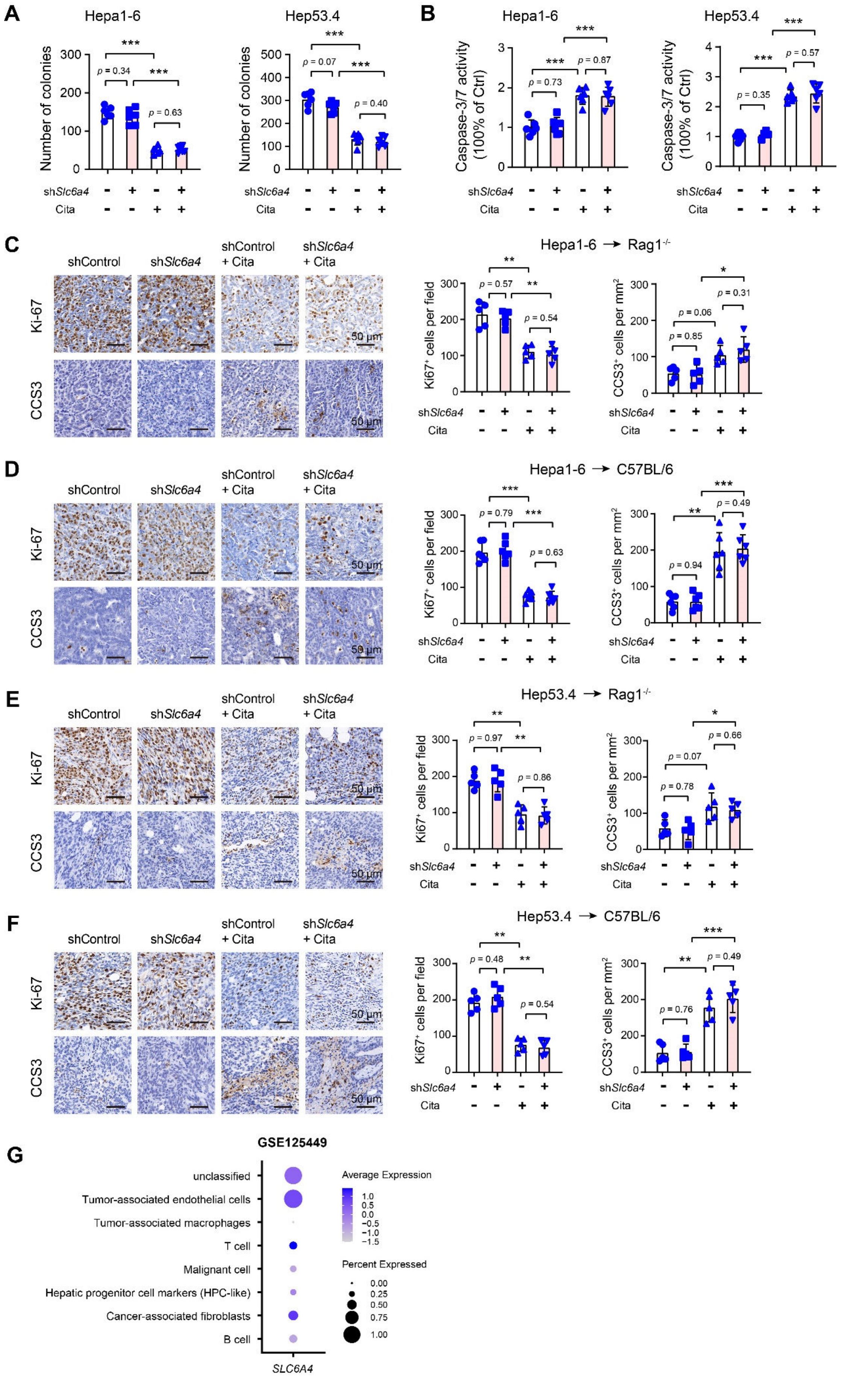

A
Hepa1-6
Hep53.4
Number of colonies
shSlc6a4
Cita
B
Caspase-3/7 activity (100% of Ctrl)
C
shControl
shSlc6a4
shControl + Cita
shSlc6a4 + Cita
Ki-67
CCS3
50 μm
Hepa1-6 → Rag1-/-
Ki67+ cells per field
CCS3+ cells per mm2
D
Hepa1-6 → C57BL/6
E
Hep53.4 → Rag1-/-
F
Hep53.4 → C57BL/6
G
GSE125449
unclassified
Tumor-associated endothelial cells
Tumor-associated macrophages
T cell
Malignant cell
Hepatic progenitor cell markers (HPC-like)
Cancer-associated fibroblasts
B cell
SLC6A4
Average Expression
Percent Expressed

**Figure S2. Citalopram suppresses HCC cell proliferation and promote cell apoptosis in a SERT-independent manner.**

(**A**) Plate colony formation assay revealed the effect of SERT knockdown alone or combined with citalopram treatment on the long-term cell proliferation of Hepa1-6 and Hep53.4 cells (n = 6 per group). (**B**) Caspase-3/7 activity in Hepa1-6 and Hep53.4 cells upon SERT knockdown or combined treatment with 5 μM citalopram (n = 6 per group). (**C**, **D**) Immunohistochemical analysis of CCS3 and Ki67 in shControl and sh*Slc6a4* Hepa1-6-bearing subcutaneous xenograft tumors from *Rag1*$^{-/-}$ or immunocompetent C57BL/6 mice, treated with DMSO or 5 mg/kg citalopram. Scale bar, 50 μm. (**E**, **F**) Immunohistochemical analysis of CCS3 and Ki67 in shControl and sh*Slc6a4* Hep53.4-bearing subcutaneous xenograft tumors from *Rag1*$^{-/-}$ or immunocompetent C57BL/6 mice, treated with DMSO or 5 mg/kg citalopram. (**G**) Single cell analysis of the expression pattern of *SLC6A4* in a HCC cohort (GSE125449). Scale bar, 50 μm. In all panels, *$p$ < 0.05, **$p$ < 0.01, ***$p$ < 0.001. Scale bar, 50 μm. Values as mean ± SD and compared by one-way ANOVA multiple comparisons with Tukey's method among groups.

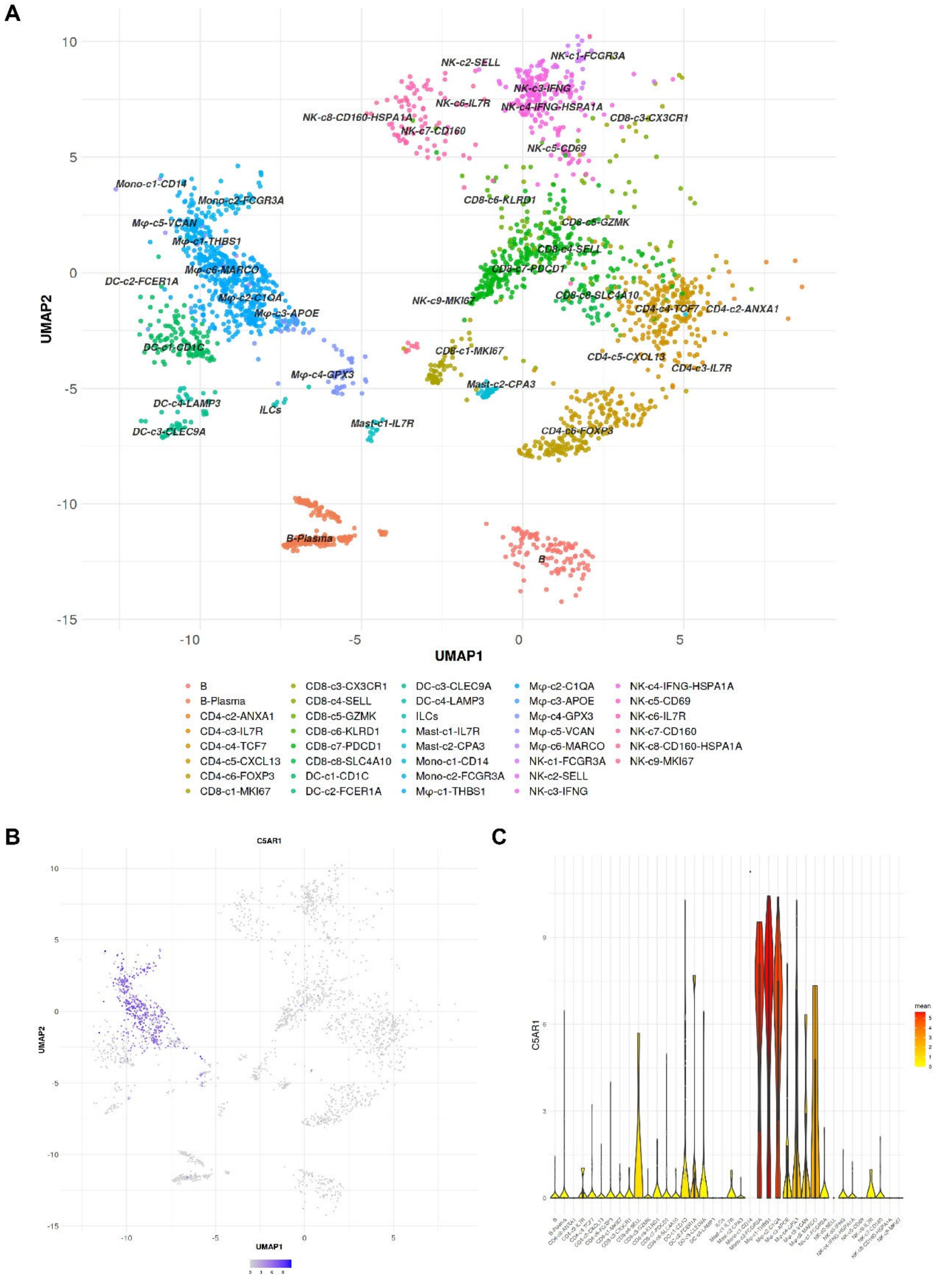


**Figure S3. The expression pattern of C5aR1 in TAMs in HCC**.

(**A**) Uniform Manifold Approximation and Projection (UMAP) of SMART-seq2-based single CD45$^+$ cells. The tSNE (by cluster) was acquired from http://cancer-pku.cn:3838/HCC/. (**B**) The UMAP showing C5AR1 expression in HCC immune cell clusters. (**C**) Violin plot showing *C5AR1* expression in different immune cell clusters.

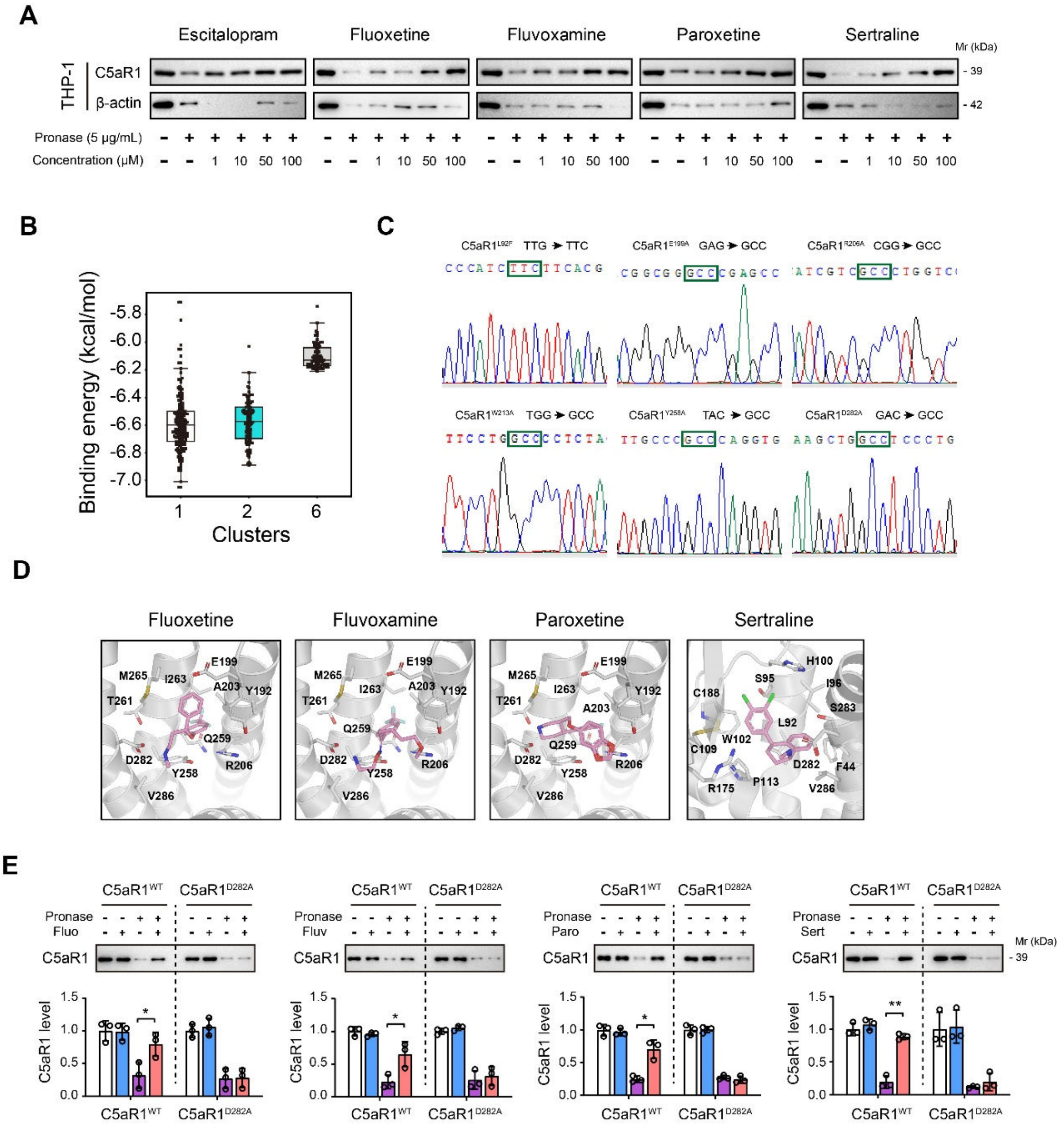

**Figure S4. Identification of C5aR1 as a direct target for citalopram**.

(**A**) The DARTS assay and immunoblot analysis showed C5aR1 protein stability against 5 μg/mL pronase in the presence of different concentrations of SSRIs treatment (0, 1, 10, 50, and 100 μM). (**B**) For C5aR1, the predicted binding energy distribution of the clusters with poses more than 50. (**C**) Sequencing analysis showed the successful generation of six C5aR1 mutants. (**D**) The best-scored complex models of C5aR1 with other four different SSRIs. (**E**) HEK293T cells were transfected with either WT or mutant C5aR1 (D282A) expression plasmids for 48 h, followed by DARTS assay with immunoblotting analysis of C5aR1 protein levels. In all panels, *$p < 0.05$, **$p < 0.01$. Values as mean ± SD and compared by one-way ANOVA multiple comparisons with Tukey's method among groups (**E**).

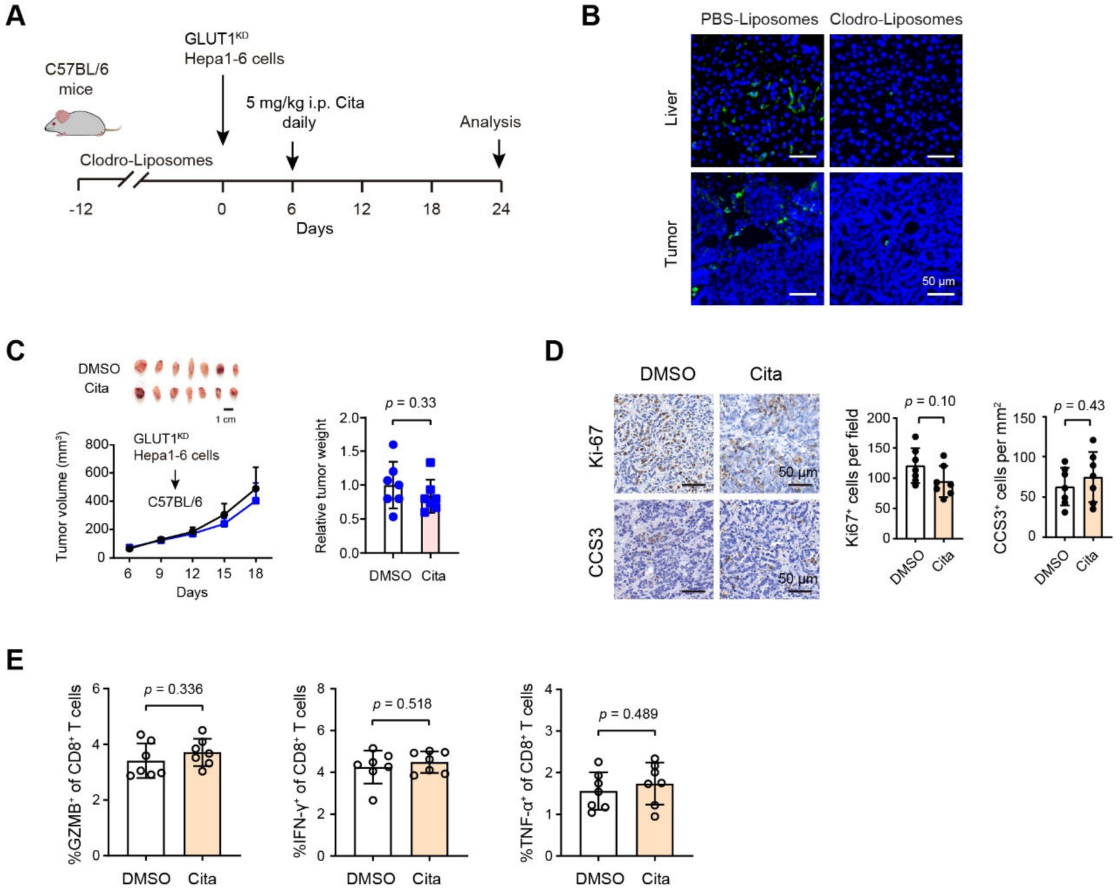


**Figure S5. Macrophage depletion mitigates the anti-tumor effects of citalopram.** (**A**) Schematic depicting macrophage blockade with clodronate liposomes. 12 days before tumor inoculation, C57BL/6 mice were pretreated with clodronate liposomes or phosphate-buffered saline (PBS) liposomes. Subsequently, GLUT1$^{KD}$ Hepa1-6 cells were subcutaneously injected into the C57BL/6 mice. The effect of citalopram (5 mg/kg) on the tumor burden was evaluated after 18 days of drug treatment. (**B**) Immunofluorescence analysis of F4/80$^{+}$ macrophages in the liver and tumor tissues of indicated groups. (**C**) In C57BL/6 mice, the effect of citalopram on the GLUT1$^{KD}$ Hepa1-6 xenograft tumors was measured in the presence of macrophage depletion (n = 7 per group). (**D**) Immunohistochemical analysis of CCS3 and Ki67 in GLUT1$^{KD}$ Hepa1-6-bearing subcutaneous xenograft tumors, treated with DMSO or 5 mg/kg citalopram (n = 7 per group). Scale bar, 50 μm. (**D**) In the context of macrophage depletion, measurement of CD8$^{+}$ T cell function in tumor tissues upon DMSO or citalopram treatment. Values as mean ± SD and compared by the Student's t test (**C**-**E**).

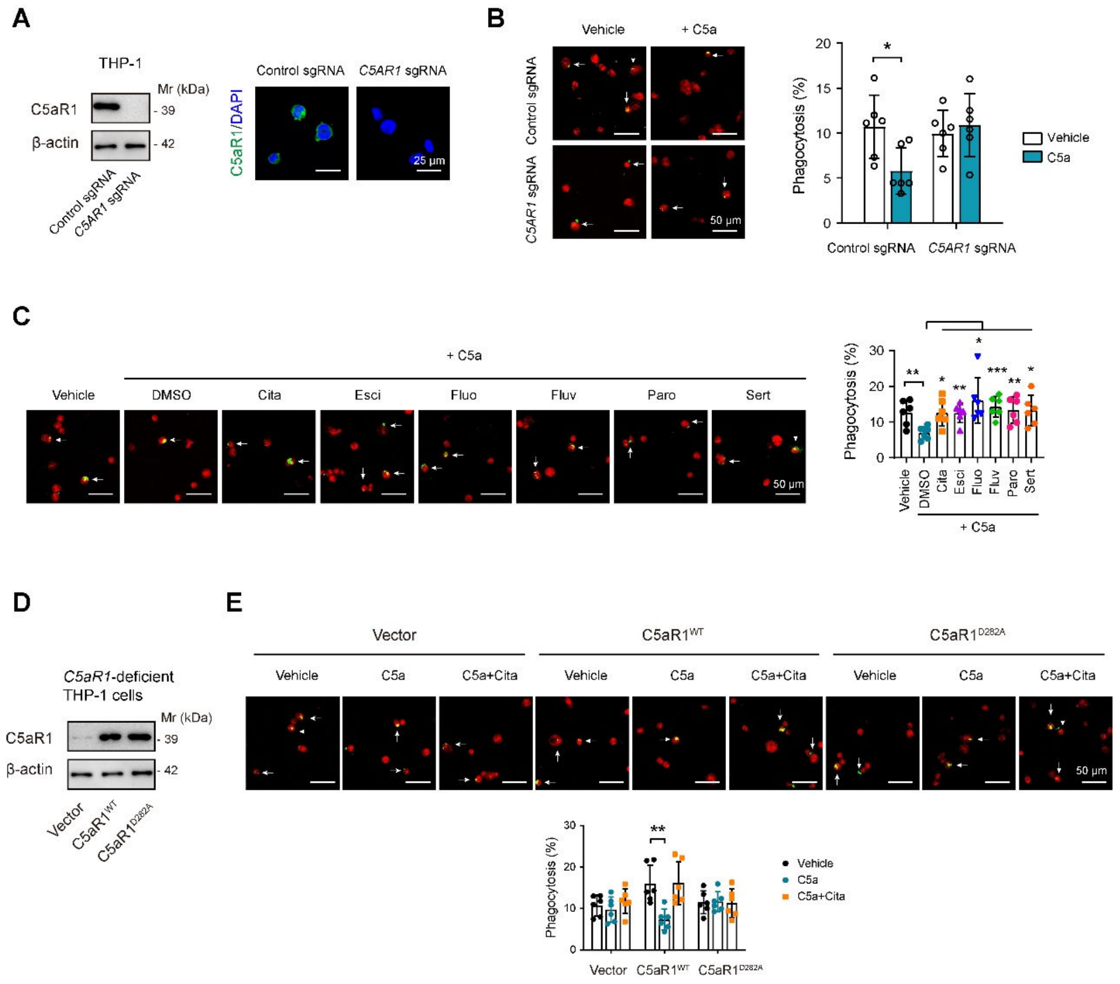


**Figure S6. Citalopram reverses C5a-mediated macrophage phagocytic impairment via targeting C5aR1.** (**A**) Western blotting and immunofluorescence analysis showed SERT protein levels in Cas9-sgControl, -sg*C5ar1* THP-1 subclones. (**B**) Effects of C5aR1 deficiency on the macrophage phagocytosis of HCC-LM3 in the presence or absence of C5a stimulation. (**C**) Effects of different SSRIs on the macrophage phagocytosis of HCC-LM3 in the presence of C5a stimulation. (**D**) Reconstituted expression of WT and D282A mutant C5aR1 in $C5aR1^{KO}$ THP-1 cells. (**E**) The effects of citalopram on macrophage phagocytosis in the absence of C5aR1 with reconstituted expression of $C5aR1^{WT}$ or $C5aR1^{D282A}$. In all panels, *$p$ < 0.05, **$p$ < 0.01, ***$p$ < 0.001. Values as mean ± SD and compared by one-way ANOVA multiple comparisons with Tukey's method among groups. Data are representative of three independent experiments (**B**, **C**, **E**).

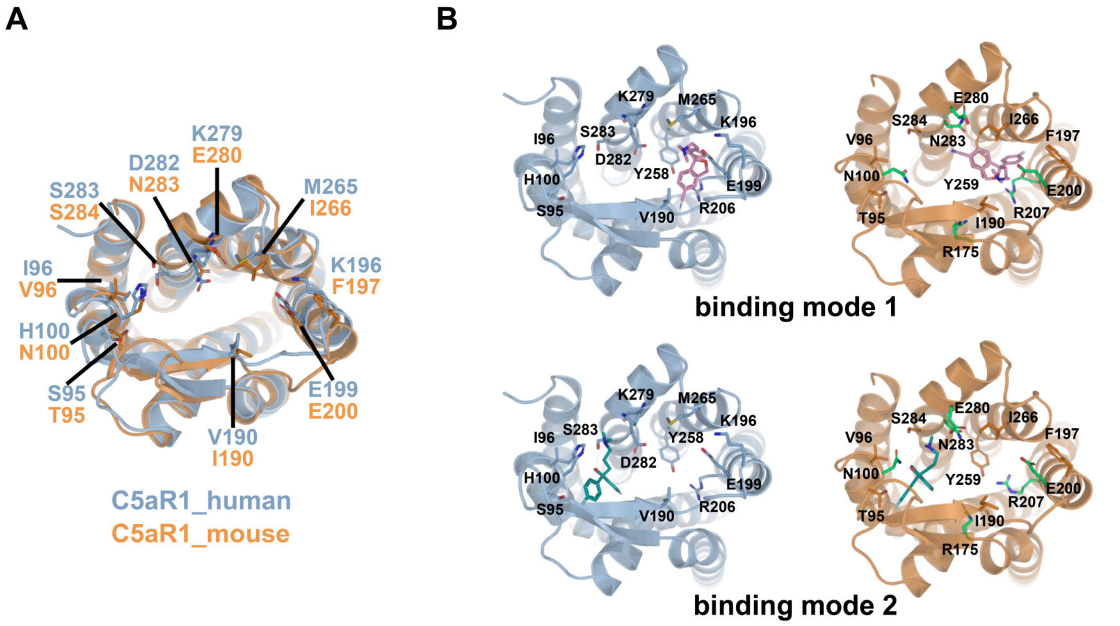


**Figure S7. Predicted binding modes of citalopram to human and mouse C5aR1**. (**A**) Conformations of orthosteric binding sites in human (light blue) and mouse (orange) C5aR1. The conformation of human C5aR1 was obtained from the crystal structure (PDB id: 6c1q). The structure of mouse C5aR1 was predicted using the ColabFold (AlphaFold2) software. (**B**) The predicted binding modes of citalopram to human (light blue) and mouse (orange) C5aR1. The conformations of citalopram were shown in pink (binding mode 1) or deep green (binding mode 2) sticks. For mouse C5aR1, green sticks indicate residues set to flexible in the molecular docking process.

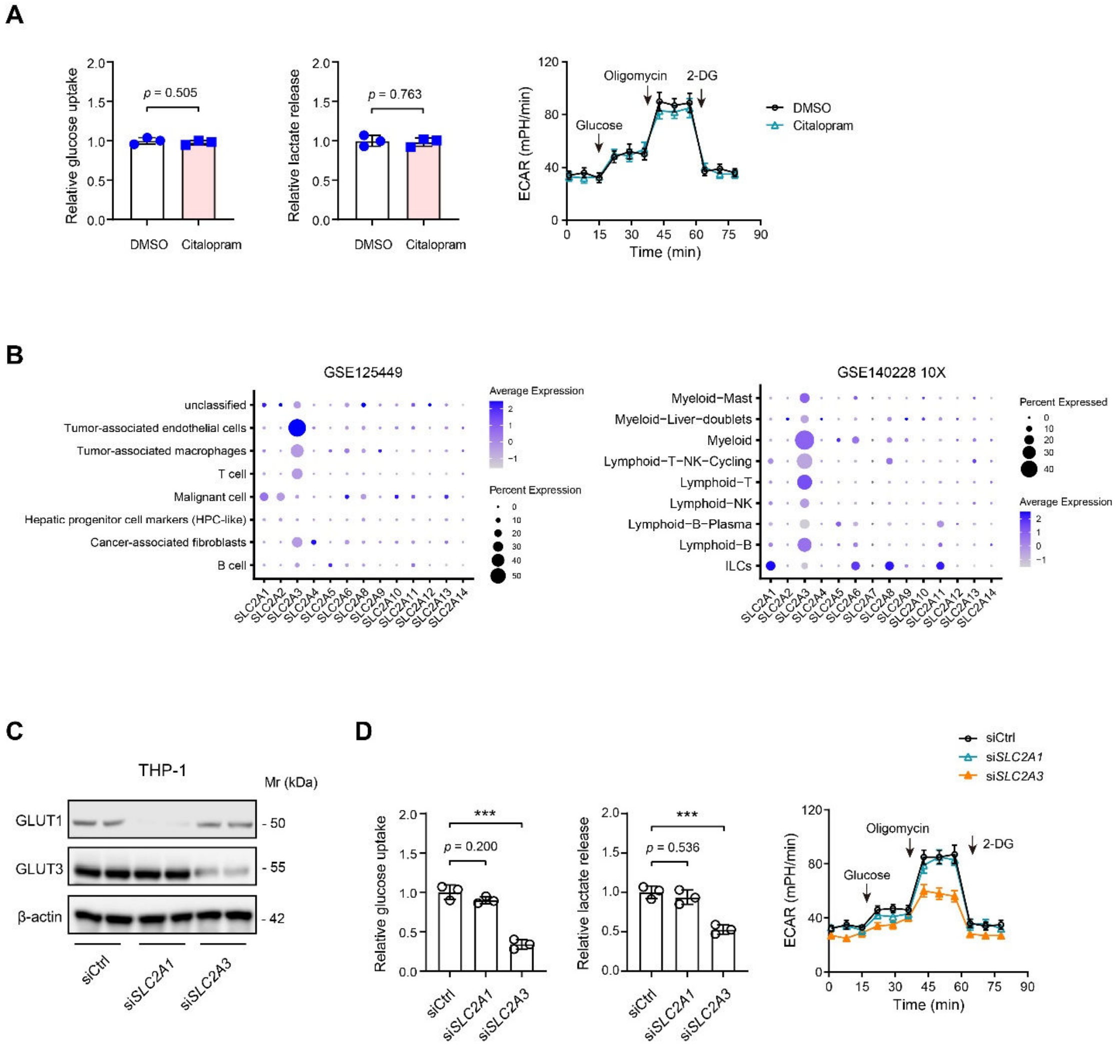


**Figure S8. Expression pattern and cellular functions of GLUT1 and GLUT3 in macrophages**.

(**A**) The effects of citalopram treatment on THP-1 cell glycolysis, as demonstrated by glucose uptake, lactate production, and extracellular acidification rate (ECAR). (**B**) Single cell analysis of the expression of glucose transporter members in two HCC cohorts. (**C**) Western blotting analysis showed the expression pattern and knockdown efficiency of GLUT1 and GLUT3 in THP-1 cells. (**D**) The effects of GLUT1 or GLUT3 knockdown on THP-1 cell glycolysis, as demonstrated by glucose uptake, lactate production, and ECAR. Values as mean ± SD and compared by the Student's t test (**A**) or one-way ANOVA multiple comparisons with Tukey's method among groups (**D**). ***$p < 0.001$.

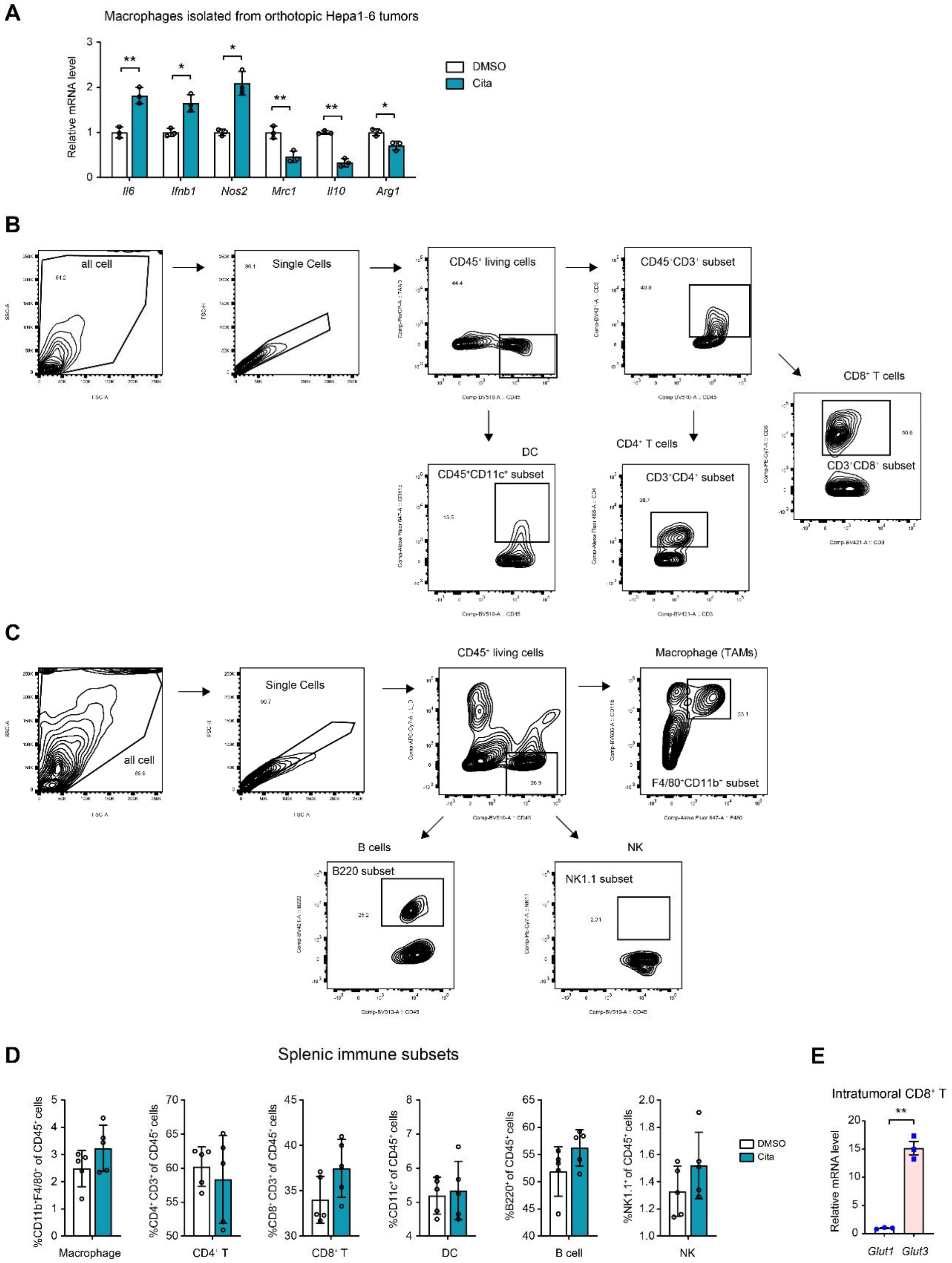


**Figure S9. Real-time qPCR analysis of macrophage polarization and flow cytometry analysis of immune cells in tumor tissues and spleen.**

(**A**) Real-time qPCR revealing the mRNA expression of M1-oriented (*Il6*, *Ifnb1*, and *Nos2*) and M2-oriented (*Mrc1*, *Il10*, and *Arg1*) markers in isolated macrophages from orthotopic Hepa1-6 tumors (n = 3 per group). (**B**, **C**) Gating strategies used for flow cytometry analysis of tumor and splenic lymphocytes. Panel A: Identification of $CD4^+$ T cells, $CD8^+$ T cells, and dendritic cells (DC). Panel B: Identification of B220$^+$ B cells, tumor-associated macrophages (TAMs), and nature killer (NK) cells. (**D**) Flow cytometry showed the infiltration of CD45$^+$CD11b$^+$F4/80$^+$ macrophages, CD4$^+$ T cells, CD8$^+$ T cells, B220$^+$ B cells, CD11c$^+$ DC cells, and NK1.1$^+$ NK cells in spleen tissues from orthotopic xenograft model, which generated in immunocompetent C57BL/6 mice with Hepa1-6 cells (n = 5 per group). (**E**) Real-time qPCR analysis of *Glut1* and *Glut3* expression in intratumoral CD8$^+$ T cells (n = 3 per group). In all panels, *$p < 0.05$, **$p < 0.01$. Values as mean ± SD and compared by the Student's t test.

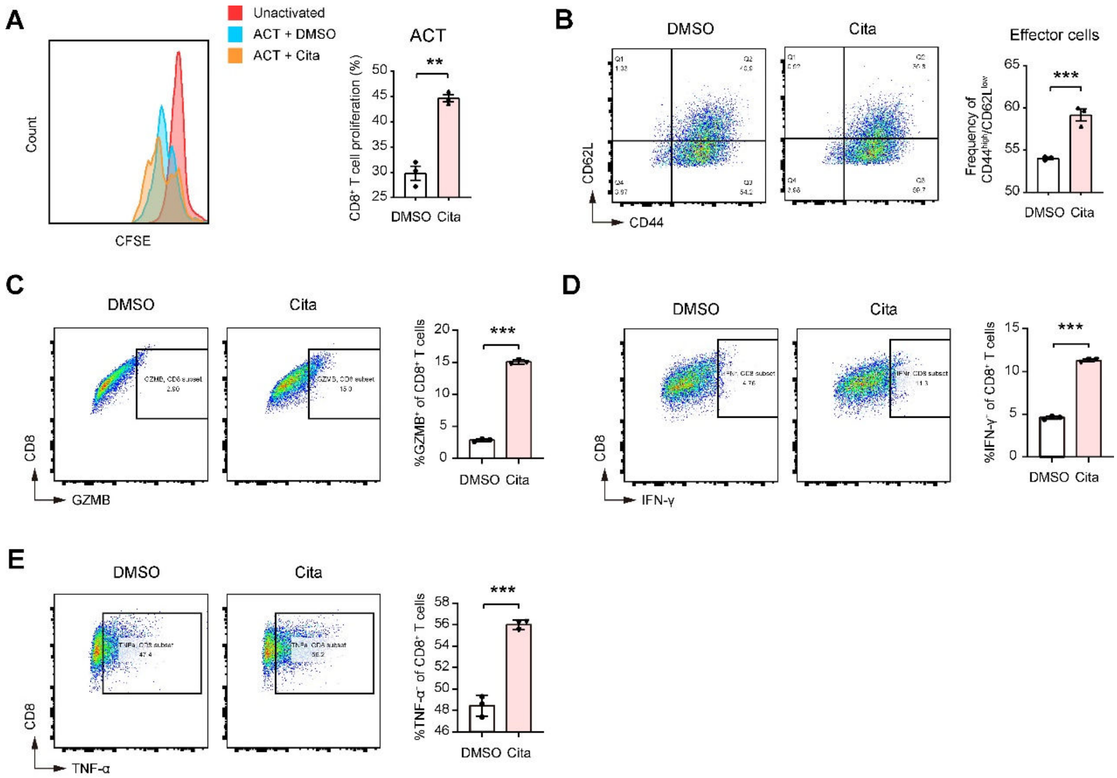


**Figure S10. The *in vitro* effects of citalopram treatment on CD8+ T cell function.**

(**A**) Splenic $CD8^+$ T cells isolated from C57BL/6 mice were stimulated with plate-bound α-CD3 plus α-CD28 for 72 h, and T-cell proliferation (CFSE staining) upon citalopram treatment was determined by flow cytometry (n = 3 per group); ACT: activated T cells. (**B**) Splenic $CD8^+$ T cells from C57BL/6 mice were stimulated with plate-bound α-CD3 and α-CD28 for 72 h, and CD44 and CD62L levels in activated $CD8^+$ T cells upon citalopram treatment were determined by flow cytometry (n = 3 per group). (**C**-**E**) Intracellular GZMB, IFN-γ, and TNF-α were detected in activated $CD8^+$ T cells upon citalopram treatment (n = 3 per group). In all panels, $^{**}p < 0.01$, $^{***}p < 0.001$. Values as mean ± SD and compared by the Student's t test.
